# IOP-induced blood-retinal barrier compromise contributes to RGC death in glaucoma

**DOI:** 10.1101/2024.10.15.618539

**Authors:** Chi Zhang, Marina Simón, Haeyn Lim, Nicholas G. Tolman, Logan Horbal, Felicia A. Juarez, Aakriti Bhandari, Christa Montgomery, Simon W.M. John

## Abstract

The integrity of the blood-retinal barrier (BRB) has been largely unexplored in glaucoma. We reveal that elevated intraocular pressure (IOP) partially compromises the BRB in two human-relevant inherited mouse models of glaucoma (DBA/2J and *Lmx1b^V265D^*). Experimentally increasing IOP in mouse eyes further confirms this. Notably, the compromise induces subtle leakage, happening without bleeding or detected endothelial cell junction disruption, and it precedes neurodegeneration. Leakage occurs from peripheral veins in the retinal ganglion cell layer with a concomitant loss of the transcytosis inhibitor MFSD2A. Importantly, stabilizing β-catenin in retinal endothelial cells prevents both vascular leakage and neurodegeneration in the DBA/2J model. The occurrence of leakage in all 3 high IOP models indicates that BRB compromise may be a common, yet overlooked, mechanism in glaucoma. These findings suggest that IOP-induced BRB compromise plays a critical role in glaucoma, offering a new therapeutic target.

## Introduction

Glaucoma is a substantial public health burden accounting for 8.4% of all blindness worldwide^1^. It currently affects ≈80 million people^2,3^, with an estimated 10% being bilaterally blind^4^. It is projected to affect >111 million people by 2040^3^. In most types of glaucoma, a harmfully high intraocular pressure (IOP) results in retinal ganglion cell (RGC) death and ultimately vision loss^5^. Multiple insults result in damage to distinct parts of RGCs in glaucoma^6,7^. In addition to direct damage to optic nerve head (ONH) axons^7–10^, there is evidence for various damaging processes within the retina, including perturbed metabolism, dendrite/synapse changes, and altered blood flow^6–8,11–22^. When developing improved treatments, it is important to understand and target distinct pathogenic events including those in the retina, ONH, and their vasculature.

The blood-retinal barrier (BRB), like the blood-brain barrier (BBB), controls and restricts the transit of molecules from blood into the surrounding neural tissue^23–25^. This maintains retinal function and prevents neural damage by immune and other harmful molecules. The BRB restricts entry of molecules into neural tissues by both maintaining paracellular endothelial cell junctions and actively suppressing transcytosis (transcellular vesicular transport)^26^. Transcytosis includes receptor-mediated and absorptive transport mechanism as well as non-specific bulk transport of fluids^27–29^. The Wnt/β-catenin signaling pathway maintains the BBB/BRB by inducing expression of tight junction proteins (e.g., claudin-5 and zonula occludens-1, ZO-1) and inhibiting transcytosis in endothelial cells to a nearly undetectable level^30–33^. Transcytosis is inhibited by inducing the expression of the major facilitator superfamily domain-containing protein 2A (MFSD2A, a key inhibitor of transcytosis). MFSD2A helps create a lipid-enriched environment that suppresses caveolar vesicle formation and transcytotic transport^32,34^. The loss of BRB or BBB integrity, either due to the loosening of tight junctions, increased transcytosis, or both triggers vascular leakage and is damaging in central nervous system diseases^26,35,36^. Despite a large volume of literature on perturbed blood flow in glaucoma, few have considered the role of the BRB in glaucoma pathogenesis^37,38^. In this study, we tested the effects of high IOP on the BRB. We investigated the BRB in three models with distinct IOP elevating mechanisms. We used two inherited, human-relevant mouse models of high IOP and glaucoma (DBA/2J and B6.*Lmx1b^V265D/+^* mice). DBA/2J mice develop a chronic form of pigmentary glaucoma caused by genes that affect melanosomal biology and are widely used for glaucoma research^39–43^. Melanosomal biology also contributes to human pigmentary glaucoma as some patients have mutations in the melanosomal gene *PMEL*^44^. Findings using DBA/2J mice have been translated to clinical trials for primary open-angle glaucoma (POAG, a common form of glaucoma), with promising initial results^13,14,45,46^. The *Lmx1b* model is directly relevant to human glaucoma as *LMX1B* is an important human glaucoma gene, contributing to both childhood glaucoma and later-onset POAG^47–50^. Here, we used mice with an *Lmx1b* mutation that causes early-onset glaucoma on the C57BL/6J mouse strain background (B6.*Lmx1b^V265D/+^*)^50^. As our third experimental system, we chose an experimentally induced model of high IOP (HAMA model) to precisely control the timing of IOP elevation and allow clear determination of the role of IOP. In this model, IOP elevation is induced via injection of a photopolymerizable, hyaluronic acid-based hydrogel into the drainage angle in front of the ocular drainage tissues^51^. This stable, biocompatible hydrogel^52^ acts as a fluid-flow resistor that elevates IOP without a need to induce tissue damage, contrary to most induced models that use lasers, saline, or beads to block and damage the drainage tissues^51,53,54^. In all three models, we show that high IOP induces BRB breakdown and vascular leakage that is detected by low molecular weight tracers but is subtle and does not involve hemorrhage. Finally, using DBA/2J mice, we experimentally rescue BRB function and show that suppressing vascular leakage reduces RGC damage, demonstrating the important contribution of vascular permeability to neurodegeneration in glaucoma and providing a new target for treatments.

## Results

### Increased IOP induces BRB compromise that precedes neurodegeneration

We first studied the BRB in the DBA/2J (D2) model. We used Hoechst tracer (MW 561.93) as it is a very sensitive marker of leakage since it binds to DNA in cells that it encounters^55^. BRB compromise detected by leakage of Hoechst tracer was present in D2 eyes that had developed high IOP. In contrast, it did not occur in any age- and strain-matched, normotensive control eyes (D2-*Gpnmb^+/+^,* hereafter called D2*-Gpnmb^+^,* which do not develop high IOP^56^) (**Figure 1A-B, Figure S1**). IOP first becomes elevated in D2 eyes at 6 months of age, with almost all eyes having high IOP by 8.5-9.5 months of age^41^. High IOP is well-established to induce corneal stretching and anterior chamber (AC) deepening in mice^42,50,57,58^. This deep-chamber phenotype confirmed exposure to high IOP in each eye with detected BRB compromise (Methods). The percentage of retinas demonstrating leakage of Hoechst tracer from retinal blood vessels in D2 eyes increased in a manner consistent with a pressure-induced etiology. Leakage occurred in 0% of D2 eyes at 6.5 months (despite some having been exposed to high IOP as evidenced by deep ACs), 30% at 7.5 months, 60% at 8.5 months, and 90% of eyes at 9.5 months of age (**Figure 1E**). The leakage was always subtle in that it was only evident by tracer monitoring with no detectable hemorrhage. Although 30% of eyes had tracer leakage at 7.5 months of age, neurodegeneration is not present at this age in D2 eyes^41^. This lack of neurodegeneration in eyes with leakage was confirmed by sensitive optic nerve analysis of >20 eyes (**Figure S2B**). Thus, BRB compromise occurs prior to neurodegeneration, raising the possibility that it contributes to glaucomatous neurodegeneration.

**Figure 1.**
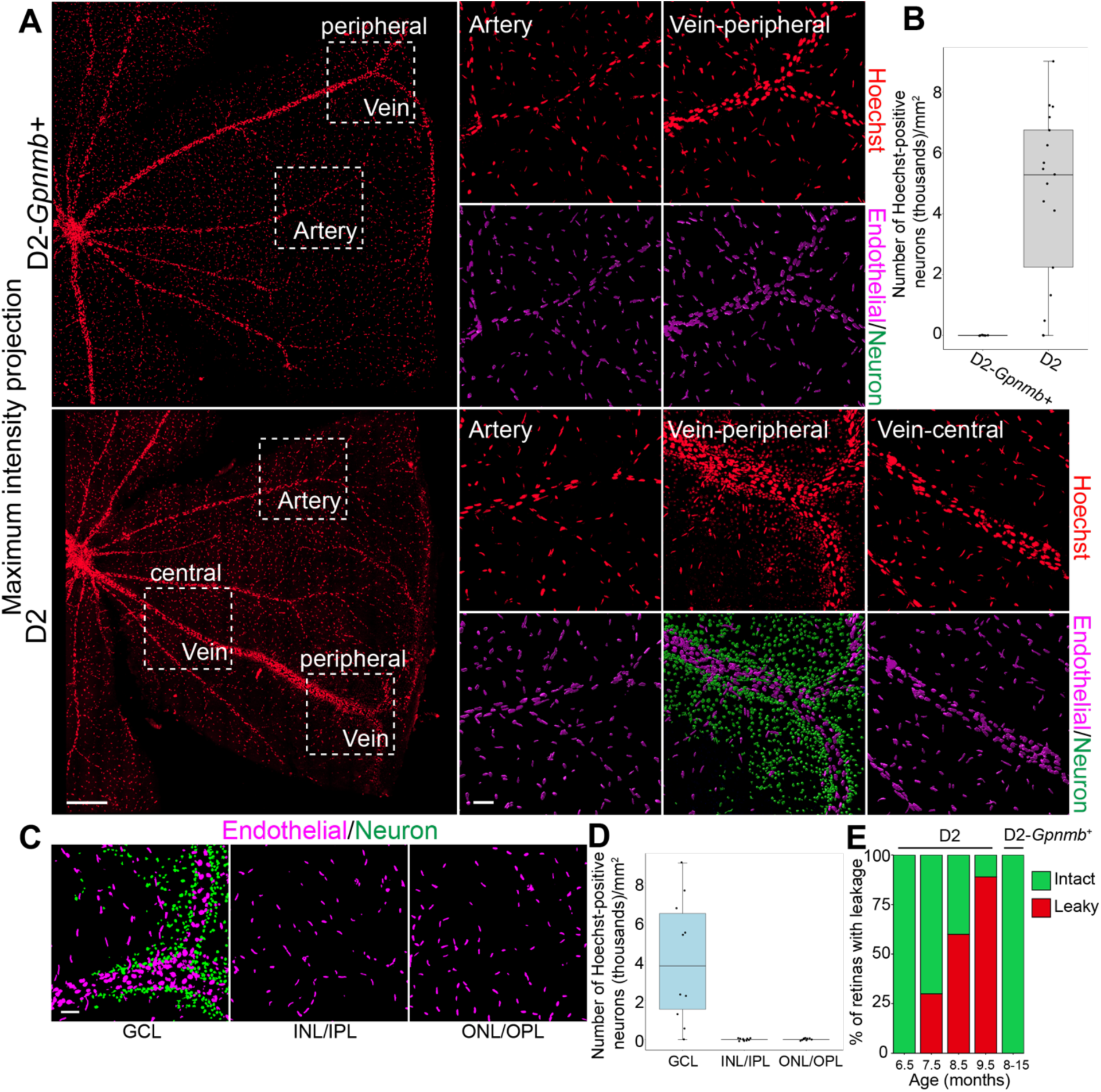
IOP-induced BRB breakdown in D2 mice. **A.** Hoechst (red) stains endothelial cell nuclei inside blood vessels throughout the retinas of control (D2-*Gpnmb^+^,* normal IOP) and D2 mice at 8.5 months of age. In ocular hypertensive D2 mice, neuronal nuclei are also stained around sites of BRB compromise where Hoechst leaks from veins. BRB compromise was initially restricted to the peripheral segments of veins as shown at their characteristic Y branch point. Imaris was used to distinguish endothelial cell nuclei (magenta) from neuronal nuclei (green) (trained machine learning function validated against cell type specific markers, see Figs. S1 and S3). Arteries were never found to leak. (Scale bar large images: 300 μm). White boxed regions are magnified on the right (scale bar: 50 μm). No leakage was detected in D2-*Gpnmb^+^*control retinas. n>20 eyes per group. **B.** As leakage only occurred from veins, the amount of leakage was quantified by counting Hoechst-positive neuronal nuclei in 0.81mm^2^ regions of the compressed z-stack that were centered over the Y branches of both peripheral veins for each flat-mounted retina. The line inside the boxplot denotes median value (50th percentile) and the box contains 70th to 25th percentiles as determined in R. Whiskers extend 1.5 times the interquartile range. D2-*Gpnmb^+^*, n = 8; D2, n = 17. **C.** Although entire retinas were studied, leakage was restricted to the veins abutting the ganglion cell layer (GCL), and no leaked tracer was detected in any retinal layer from inner nuclear layer (INL) to outer nuclear layers (ONL) in 8.5-month-old D2 mice. Scale bar: 50 μm. n = 40 eyes. **D**. Quantification of leakage in GCL, INL/IPL, and OPL/ONL with the same boxed region quantified at different tissue depths corresponding to each retinal layer. n = 10. **E.** Percentage of retinas with leakage across age in D2 mice. n > 20 for each age. IOP changes with age are well established in D2 mice at these ages (see ref 120). No D2 mice had leakage at 6.5 months of age, when IOP first becomes elevated. By 8 to 9 months of age, most D2 eyes have high IOP. Increased anterior chamber depth confirmed exposure to high IOP in all eyes with BRB compromise. These data show that IOP elevation precedes BRB leakage.

### In D2 mice, detected leakage occurs from veins and is restricted to the ganglion cell layer

Hoechst tracer that leaks from blood vessels initially stains nuclei adjacent to the leak. This allowed us to precisely locate sites of BRB compromise. Hoechst-stained neurons are first detected next to the peripheral segments of veins, where they branch in a Y shape as they travel around the far periphery of the retina (**Figure 1A**). With longer exposure to high IOP and increased leakage, tracer spreads further across the tissue labeling neurons distant from vessels and throughout the ganglion cell layer (GCL) (in some by 9 months and all by 9.5 months of age). No leakage was ever observed arising from arteries or capillaries. Importantly, leaked tracer was restricted to the GCL, consistent with leakage from the major retinal veins of the inner retina and not capillaries in other retinal layers (**Figure 1C, D**). No leaked tracer was present in the retinal tissue extending from the inner to outer nuclear layers even with more extensive aging to 14 months of age (**Figure S3)**.

### IOP-induced BRB compromise appears to be a general feature of glaucoma

IOP elevation in D2 mice is caused by a pigment-dispersing iris disease that induces a pigmentary form of glaucoma^40,41,59^. To determine if IOP-induced BRB compromise occurs in eyes with a very different IOP elevating etiology and to initially assess its relevance to more common human glaucoma, we next tested the BRB integrity in mice with a mutation in a human glaucoma gene, *Lmx1b,* which contributes to a spectrum of human glaucoma including POAG^47–50^ (**Figure S4**). B6.*Lmx1b^V265D/+^* mice develop high IOP beginning at 2 months of age (**Figure S6**) and have glaucomatous degeneration starting at 6.5 months of age^50^. Leakage of Hoechst tracer again indicated BRB compromise in ocular hypertensive *Lmx1b^V265D/+^* mice but not in normotensive *Lmx1b^+/+^* control littermates (**Figure S4A-B, Figure S5**). Tracer leakage was evident in 5.5-month-old eyes prior to neurodegeneration. As with D2 mice, BRB compromise appeared venous-specific, with tracer leakage restricted to the GCL (**Figure S4E-F**).

As a final model with another distinct IOP elevating etiology, we induced IOP elevation using a photo-polymerizable HAMA hydrogel (HAMA)^51^. Although not a genetic model with a human counterpart, this model allowed us to definitively control the timing of IOP elevation and assess the effects of high IOP without the confounding effects of other ongoing disease processes. Placing this hydrogel, fluid-flow resistor in front of the ocular drainage tissues reproducibly elevates IOP without requiring tissue-damaging/blocking processes that are typically more variable. This model reproducibly induces a sustained elevation of IOP (**Figure S7**). To determine the effects of experimentally elevating IOP on the BRB, we compared ocular hypertensive C57BL/6J eyes that had polymerized HAMA (pHAMA) to normotensive strain-matched control littermates that were administered HAMA but without photopolymerization (npHAMA. **Figure S4C-D, Figure S7**). BRB leakage was detected in ocular hypertensive but not normotensive eyes. Again, BRB compromise preceded neurodegeneration (**Figure S3**), was venous-specific, and was restricted to the GCL (**Figure S4G, H**). The documented presence of BRB compromise in ocular hypertensive eyes of mice with two distinct types of inherited glaucoma on two genetically distinct strain backgrounds as well as following experimentally elevated IOP suggests that high IOP may generally induce BRB compromise.

### Tight junctions and mural cells appear normal

Since they are a widely used model of chronic glaucoma with findings that have translated to human glaucoma^13,14,45,46^, we continued to use ocular hypertensive D2 mice to investigate the mechanisms of BRB compromise. The BRB is maintained by the neurovascular unit, including pericytes and endothelial cells, with a key component being tight junctions (TJs) between endothelial cells^60^. Mural cells including pericytes are key regulators of the BRB by acting to limit transcytosis and modulate tight junction formation^61^. BRB breakdown can result from compromised TJs or loss/dysfunction of mural cells^62^. Thus, we investigated the integrity of TJs as well as the distribution and morphology of mural cells/pericytes by immunofluorescence in retinal vessels of D2 and D2-*Gpnmb^+^*mice. No differences in fluorescence intensity of TJ components or mural cells/pericytes were detected between ocular hypertensive D2 and normotensive control mice (**Figure 2**). The expression levels of TJ proteins claudin-5, ZO-1, and occludin were also unchanged when assessing whole retinal lysates by Western blotting in D2 mice (**Figure 2E-F and Figure S10**). Finally, we analyzed claudin-5 staining in retinal flat mounts of pHAMA and B6.*Lmx1b^V265D/+^* mice via immunofluorescence and confirmed that there was no difference in fluorescence intensity or distribution pattern between the retinas of ocular hypertensive mice and their controls **(Figure S8**). These findings suggest that neither TJ composition nor mural cell/pericyte loss are the main drivers of BRB disruption after IOP elevation.

**Figure 2.**
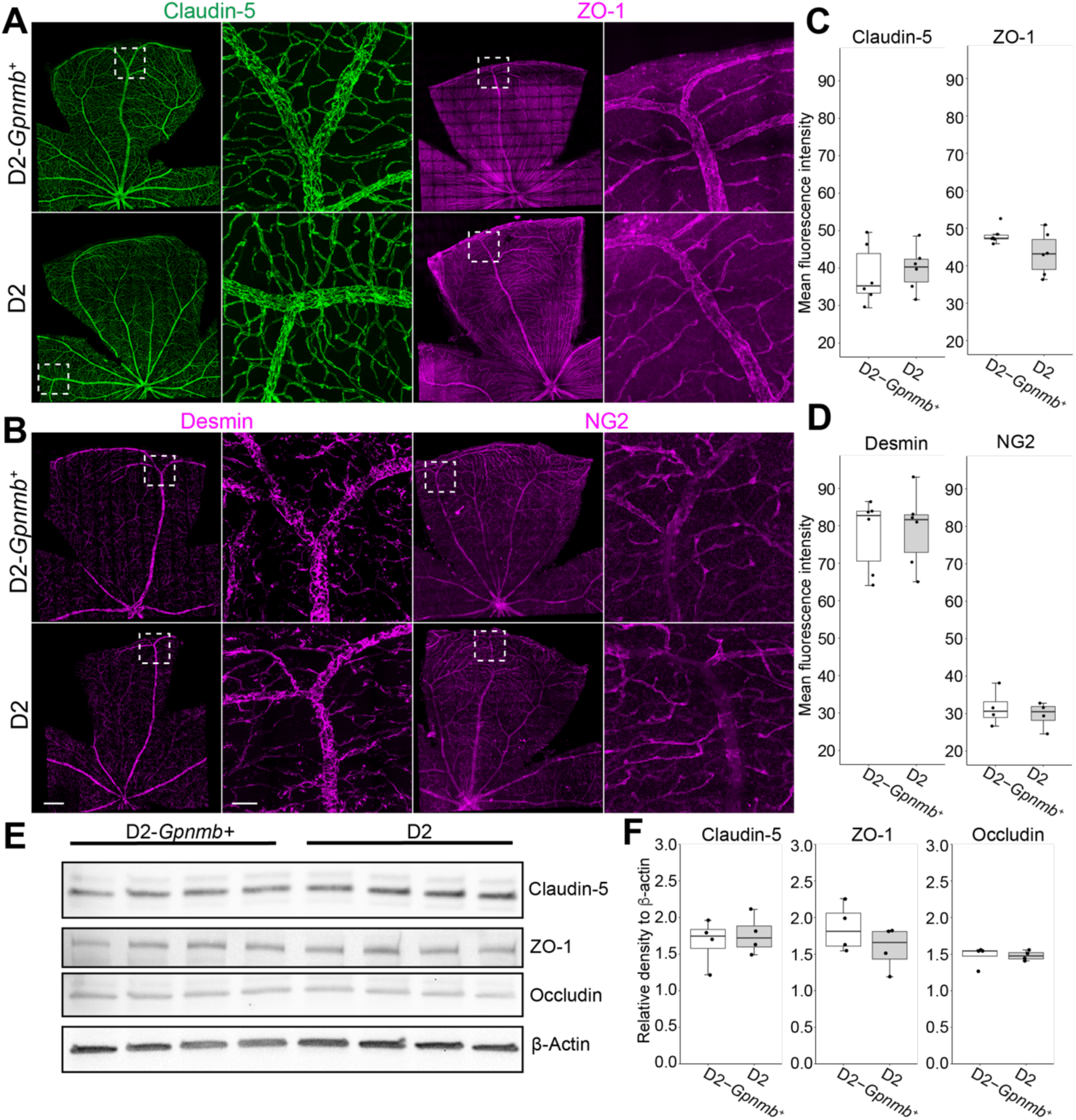
Normal appearing tight junctions (TJ) and pericytes in the retinal vasculature of D2 mice. A-B. Representative images of tight junction (A, claudin-5 and ZO-1) and pericyte markers (B, desmin and NG2) in retinal flat mounts of 9-month-old normotensive control (D2-*Gpnmb^+^*) and ocular hypertensive D2 mice. No differences were detected. Paired images for each marker with boxed region magnified in the right image of each pair. Scale bars: left images, 300 μm; right images, 50 μm. **C-D.** Quantitative analysis of fluorescence intensity in peripheral veins (Methods) by measuring mean value in Image J. The fluorescent signal was not significantly different between D2-*Gpnmb^+^*and D2 mice for any of the markers analyzed (means for each marker were compared using the Welch’s t-test. *P* = 0.70 for claudin-5, *P* = 0.095 for ZO-1, *P* = 0.82 for desmin, and *P* = 0.55 for NG2). n = 6 veins each group for all except n = 4 veins each group for NG2. **E-F.** Western blot analysis of 15-month-old retinas from eyes with confirmed exposure to high IOP. Even after more extensive IOP/glaucoma exposure at older ages, no differences in the expression levels of TJ proteins ZO-1, occludin, and claudin-5 were observed. The density relative to β-actin was quantified for each protein. (Means for each protein were compared using the Welch t test. *P* = 0.6726 for claudin-5, *P* = 0.2644 for ZO-1, and *P* = 0.9822 for occludin. n = 4 each group).

### Abnormally active transcytosis may underlie IOP-induced BRB compromise

The lack of detected changes in tight junction composition and abundance led us to investigate a transcytotic mechanism of BRB disruption. Albumin transport is generally considered a marker of transcytosis^63^. Thus, we compared albumin transport across the vascular endothelium in D2 and normotensive control mice (**Figure 3**). Albumin remained confined within retinal vessels in age- and sex-matched normotensive D2-*Gpnmb*^+^ mice, reflecting an intact BRB with strongly repressed transcytosis. In contrast, albumin leakage was readily evident from peripheral retinal veins (but not arteries or capillaries) in ocular hypertensive D2 mice of both sexes, consistent with excessive transcytosis due to deficient repression of vesicular transport. Importantly, albumin extravasation was also confined to the GCL with no leakage detected from deeper retinal vascular plexuses (**Figure 3C**). This suggests that abnormally active transcytosis contributes to IOP- induced BRB compromise.

**Figure 3.**
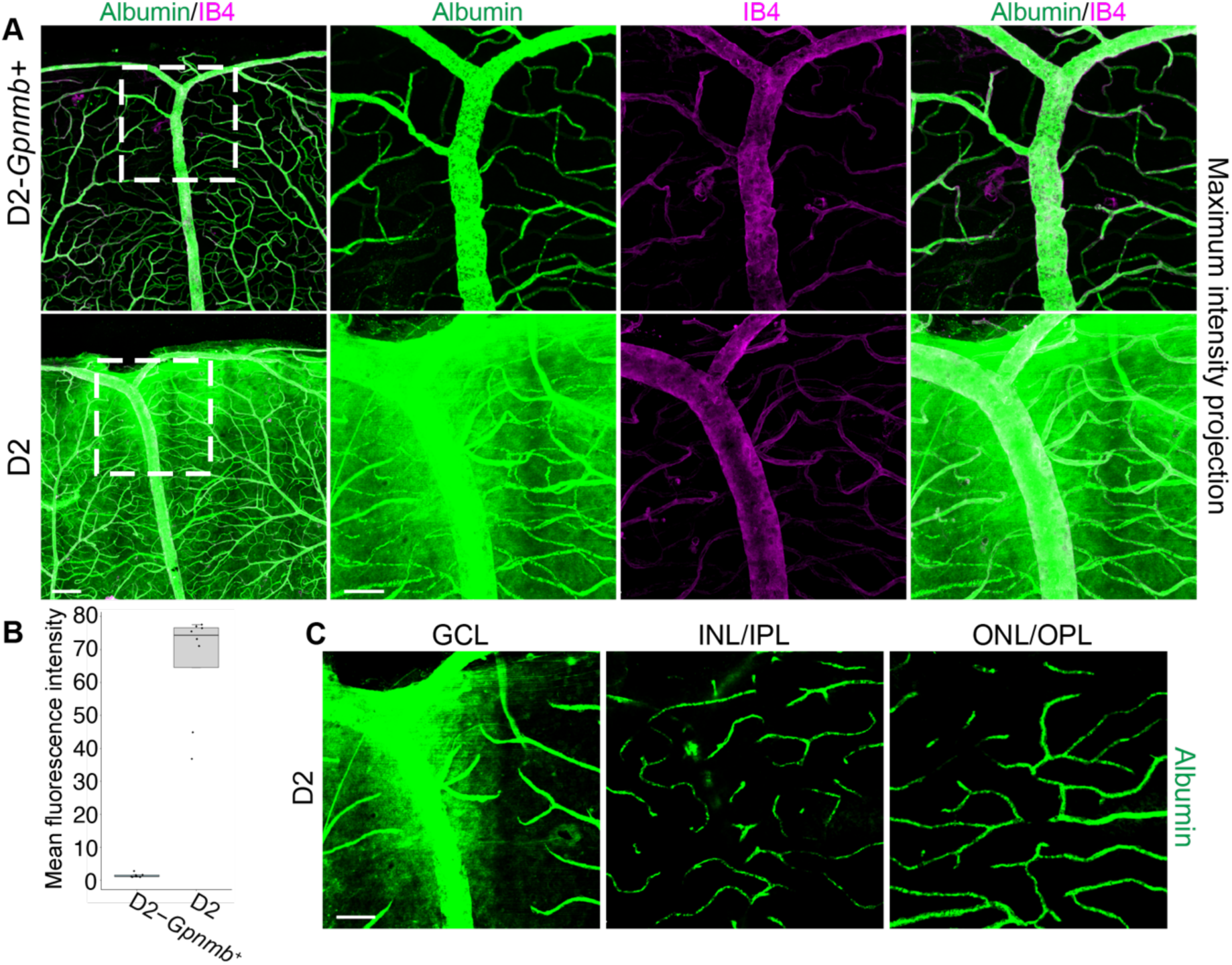
Albumin leakage from D2 retinal veins. **A.** Albumin (green) and IB4 (magenta, blood vessel marker) immunostaining in retinal flat mounts of 9-month-old D2-*Gpnmb^+^*and D2 mice. The dashed boxes are magnified in the adjacent images for albumin alone, CD31 alone, and both markers merged (rightmost panels). Albumin is constrained inside the vasculature in normotensive D2-*Gpnmb^+^* eyes but leaks into the retinal parenchyma in hypertensive D2 eyes. Scale bars: leftmost images, 300 μm; magnified images, 50 μm. **B**. Quantification of extravasated albumin (See methods) across the entire retina as mean fluorescence intensity. n = 8 eyes per group. **C**. Albumin leakage is restricted to the GCL layer in D2 mice. n = 8 eyes. Representative images are of the same tissue location at different depths, while quantifications were performed on whole retinas.

Given the role of MFSD2A as a key transcytosis inhibitor in CNS veins and capillaries (but not arteries^64,65^), we next investigated if it is affected by IOP (**Figure 4**). Further supporting a transcytotic mechanism of barrier compromise, MFSD2A immunofluorescence was decreased in endothelial cells of peripheral retinal veins of ocular hypertensive mice. In contrast, MFSD2A immunofluorescence was not altered in capillaries of the same hypertensive eyes or in any vessels of normotensive control eyes (**Figure 4A**). Furthermore, the regional loss of MFSD2A matched the locations of BRB compromise (**Figure 4B**). For example, albumin leaked at sites with no detectable MFSD2A but was constrained to the vascular lumen in regions where endothelial cells still expressed MFSD2A, even if that MFSD2A expression was at lower-than-normal levels. These data argue that excessive transcytosis due to loss of MFSD2A may be the cause of BRB dysfunction following IOP elevation in D2 mice.

**Figure 4.**
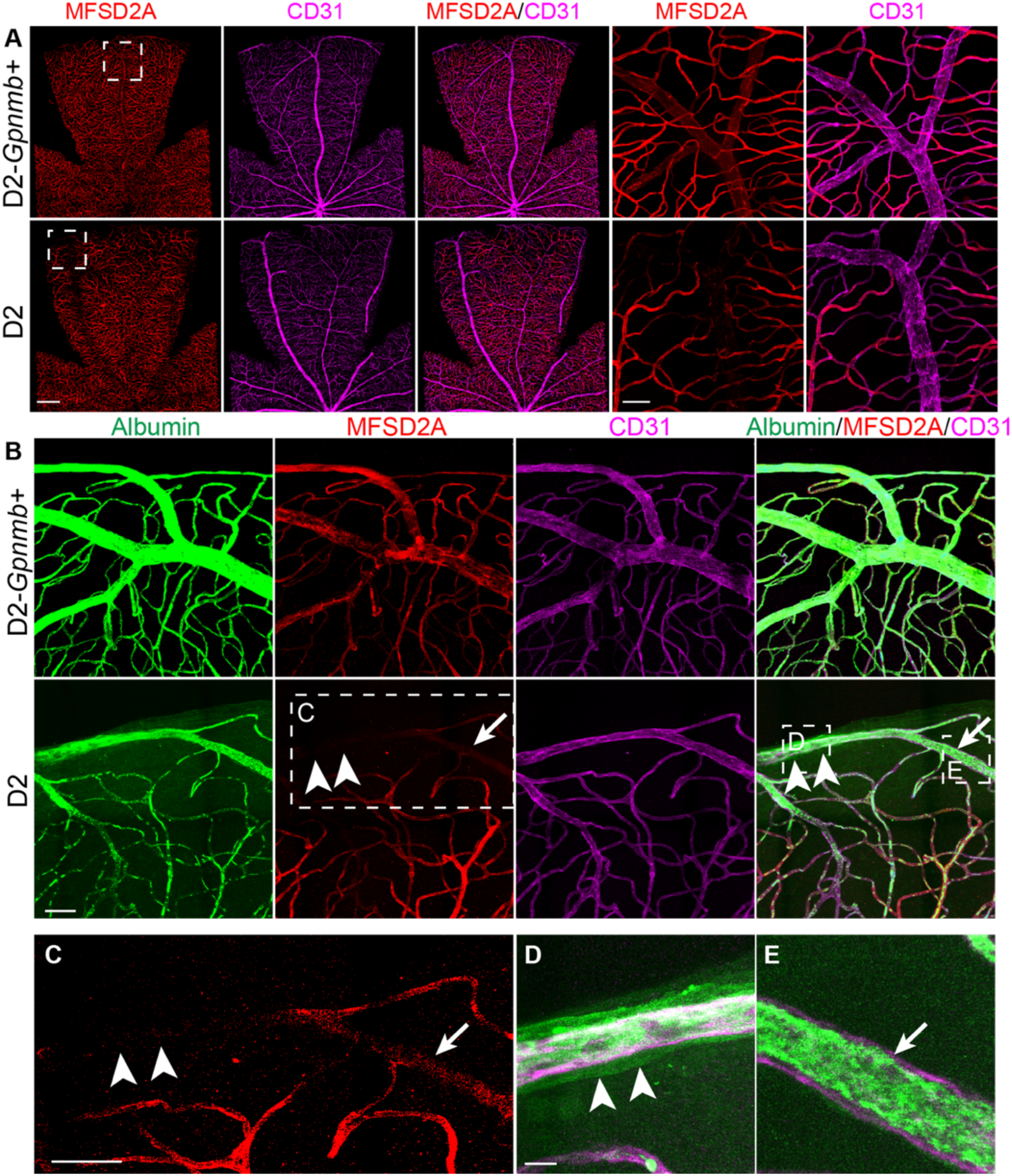
MFSD2A expression is lost in retinal veins of D2 mice and colocalizes with albumin leakage. **A**. MFSD2A (red) and CD31 (magenta, endothelial marker) expression in retinal flat mounts (All 9-month-old). Dashed boxes are magnified in the two right-most panels. MFSD2A is decreased or lost in retinal veins of D2 eyes that had high IOP, but not in normotensive D2-*Gpnmb^+^* eyes. Scale bars: first 3 panels, 300 μm; magnified images, 50 μm. **B.** Albumin leaks from veins at sites with severe (undetectable, double arrowheads) but not partial (arrow) loss of MFSD2A. Scale Bars = 50 μm. **C-E.** Magnified images of corresponding white dashed boxes in B show this more clearly. Albumin is contained inside the vessel at the site with reduced MFSD2A (E), but albumin has leaked from the vascular lumen and coats the vessel wall where MFSD2A is not detectable (D). All 9-month-old. n = 8 veins/4 eyes each group (or 8 venous segments without MFSD2A, 8 with decreased MFSD2A, and 8 control veins). Scale bars: C, 50 μm; D and E, 10 μm.

### Stabilization of β-catenin prevents BRB breakdown and ameliorates glaucoma

To test if the BRB leakage damages RGCs in glaucoma, we asked if specifically stabilizing β-catenin in endothelial cells rescues the BRB and modulates glaucoma development. We used an endothelial cell-specific Cdh5-Cre/ERT2^66^ to conditionally activate the expression of a stabilized allele of β-catenin (*Ctnnb1^flex3^*) (**Figure 5**). This allele encodes a form of β-catenin that cannot undergo normal phosphorylation and so is degraded at a decreased rate^67^. Tamoxifen was administered to 6.5-month-old D2 mice to activate the Cre, which in turn activated the stabilized β-catenin allele (D2.*Ctnnb1^flex3^*^/+^; Cdh5-Cre/ERT2 mice). Vascular endothelial β-catenin stabilization did not alter the IOP-elevating iris disease or IOP itself (**Figure S9**). However, the stabilized β-catenin ameliorated vascular leakage (**Figure 5A-B**) and decreased glaucomatous optic nerve and retinal damage (**Figure 5C-E**), compared to control mice (tamoxifen injected D2.*Ctnnb1^flex3^*^/+^ mice). Thus, β-catenin stabilization restored BRB function and significantly lessened glaucoma in D2 mice.

**Figure 5.**
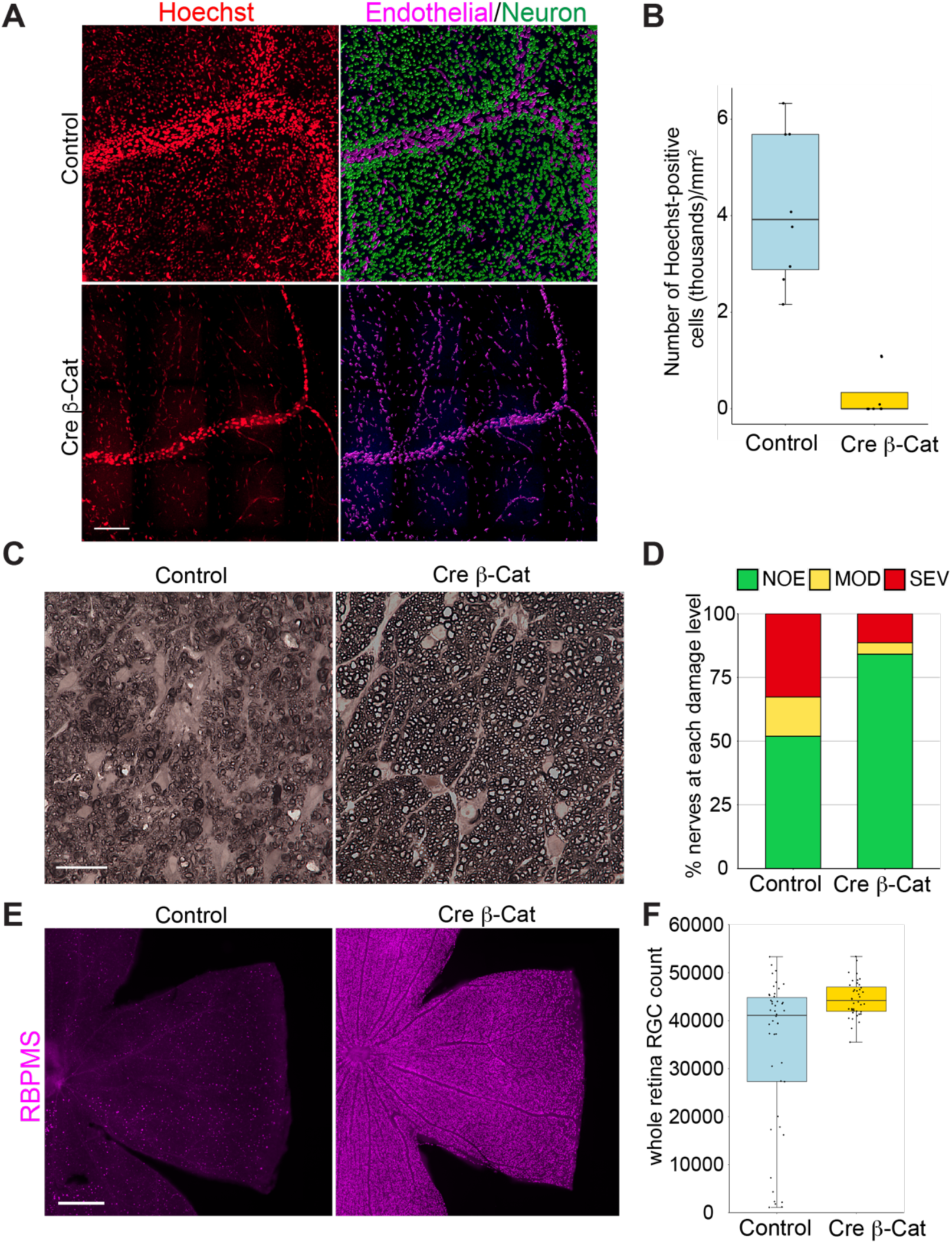
β-catenin stabilization restores BRB function in D2 mice. **A.** Hoechst staining (red) in 9-month-old D2.*Ctnnb1^flex3^*^/+^ (Control) and D2.*Ctnnb1^flex3^*^/+^; Cdh5-CreERT2 (Cre β-Cat) mice. Machine-learning filtered images (last panel) differentiating Hoechst-positive endothelial cells (magenta) vs. neural cells (green) demonstrate that expression of stabilized β-catenin stops vascular leakage in D2 mice. Scale bar: 100 μm. n>20 eyes each group. **B.** Quantification of leakage. Leakage was significantly reduced in the stabilized β-catenin-expressing group (Welch’s t-test, *P* = 0.00012; n = 8 each group). **C.** Representative images of optic nerves. Control eyes that do not express stabilized β-catenin develop severe glaucomatous nerve damage. The majority of Cre β-Cat eyes have no nerve damage. Scale bar: 50 μm. **D.** Nerve damage was significantly reduced in Cre β-Cat mice at 9 months (Fisher’s exact test, *P* = 0.0035). SEV = severe (>50% RGC axon loss), MOD = moderate (average 30% axon loss), NOE = no glaucomatous damage (undetectable axon loss). n = 52, control; 44, Cre β-Cat. **E.** Representative images of RBPMS stained retinas showing preservation of RGCs in Cre β-Cat retinas at 9 months. Scale bar: 500 μm. **F**. RBPMS stained RGC counts (entire ganglion cell layer) were significantly elevated by β-catenin stabilization in Cre β-Cat mice vs. Control mice (Welch’s t-test, *P* = 8.051e-05. Control, n = 46; Cre β-Cat, n = 40.)

## Discussion

Here, we document an IOP-induced compromise of the BRB in retinal veins that colocalizes with loss of MFSD2A. This compromise allows low molecular weight molecules and proteins, but not cells, to leak from veins into the retinal parenchyma. Further, we experimentally determine that this subtle compromise precedes RGC loss and contributes to glaucoma progression. By specifically stabilizing β-catenin within vascular endothelial cells, we prevent leakage and ameliorate RGC death. Vascular leakage occurred following IOP elevation of three distinct etiologies and on different genetic backgrounds. This suggests that BRB compromise may be a general feature driving glaucoma. Recently, elevated blood pressure was shown to cause focal vascular leakage with a similar decrease of MFSD2A in the brains of spontaneously hypertensive rats^68^. Together with our data, this supports pressure-based modulation of MFSD2A in different species. Interestingly, mutations in some genes that contribute to POAG (*CAV1, FOXC1*) have been shown to disturb the BRB/BBB in animal studies^26,37,64,69–71^.

In all three ocular hypertensive models studied here, BRB leakage was only detected in the GCL. In other studies of vascular leakage caused by β-catenin signaling pathway mutations (e.g., LRP-5, Frizzled-4), leakage occurs from both veins and capillaries of all retinal layers^72,73^. This difference is likely related to those mutations profoundly affecting signaling in all of these vascular beds. For unknown reasons (possibly mechanical), IOP elevation in our models was only detected to affect the peripheral segments of major veins at the inner retinal surface. Even in 9.5-month-old D2 eyes, when all GCL neurons are labeled by Hoechst, albumin leakage was only detected from peripheral veins abutting the GCL (most albumin detected on and around vein). This GCL-specific pattern of leakage in the ocular hypertensive models may be another factor contributing to the unique vulnerability of RGCs versus other retinal neurons to glaucoma. Different insults damage different parts of the RGC, with a key insult damaging RGC axons in the ONH^7–10^. This axonal insult is believed to be a critical insult for RGC demise, as the topography of RGC dysfunction/loss and the pattern of visual field deficits are consistent with the bundling of axons in the ONH^6,7,9,74,75^. The generally accepted necessity of the ONH insult and sparing of amacrine cells (another major neuronal population in the GCL) may suggest that multiple hits are necessary to kill RGCs, with the partial BRB compromise alone being insufficient. Additionally, amacrine cells may be more resilient than RGCs to the local environmental changes following IOP elevation and BRB compromise. This is supported by the greater vulnerability of RGCs in both diabetes models and Alzheimer’s disease, conditions with vascular leakage in the retina^76–78^.

Our data suggest that increased transcytosis due to loss of MFSD2A, but not TJ defects, is the main driver of BRB compromise in D2 mice. Our existing data support the same mechanism in the other tested models of IOP elevation. Previous studies have convincingly demonstrated that experimental ablation of *Mfsd2a* in mice results in BBB/BRB leakiness due to increased vesicular trafficking across endothelial cells^30,64,79^. However, other studies have not observed an effect of MFSD2A mutation on vascular leakage^80–82^. Overall, this reveals complexity that is not yet understood and may be related to genetic, experimental and environmental differences. Our data further support a role of MFSD2A in regulating the BRB and transcytosis, but further work is needed to definitively demonstrate a transcytotic mechanism in glaucoma. Consistent with our data in D2 mice, previous studies have shown that experimentally stabilizing β-catenin/Wnt signaling rescues vascular barrier defects in different conditions, including in Alzheimer’s disease and diabetic retinopathy models^76,77^.

The exact mechanisms by which high IOP induces endothelial cell changes and BRB breakdown remain to be elucidated. Studies have shown that high IOP leads to oxidative stress and other cellular stresses (that mimic or exacerbate age-dependent changes), causing oxidative damage and metabolic disturbances to cells^83^. Stress and metabolic alterations in endothelial cells, mural cells, or other neurovascular unit cells would likely result in BRB deterioration over time. Alternatively, signaling by mechanosensing molecules in endothelium may be modulated by IOP in a way that affects barrier integrity^84^. Mechanosensing molecules include VE-Cadherin, and integrins^84^, but they are not known to modulate MFSD2A. Understanding these mechanisms will identify new therapeutic targets, potentially transforming the treatment of glaucoma and other diseases characterized by BRB dysfunction.

BRB compromise, similar to that reported here, may occur commonly in POAG and other forms of human glaucoma. It could have been previously missed as tracers need to be used for detection, and the leakage could be restricted to the far peripheral regions of retinal veins. In the clinic, glaucoma patients are not normally evaluated using vascular tracers, and the vasculature in the far retinal periphery is rarely assessed even in research studies. Inflammation is known to compromise vascular barriers and is clinically known to cause more generalized vascular leakage in a subset of eyes with inflammatory conditions and glaucoma^85,86^. We previously reported leakage in D2 mice but these initial experiments neither determined the peripheral venous or molecular nature of leakage nor tested the causative role of IOP^87^. In the *Lmx1b* mouse model, some leakage also occurred from vessels in the ONH, but further experiments are needed to determine both how common this is and the timing of its onset in relation to retinal vein leakage. Importantly, studies in humans have reported subtle vascular leakage (tracer leakage without hemorrhage) in the ONH of patients with POAG^88–101^. The leakage was present in 30% or more of POAG patients^93^ and was significantly correlated to IOP^101^. Despite this and an extensive literature on the vasculature and blood flow^102–104^, BRB compromise has hardly been considered in glaucoma. Neither IOP-induced BRB compromise nor vascular leakage were previously shown to contribute to glaucoma progression.

BRB disruption occurs in Alzheimer’s disease and other neurodegenerative diseases^105–109^, but is less specific and more severe than we document here in glaucoma. In Alzheimer’s disease, it disrupts tight junctions and affects the extensive capillary network in the retina^110–112^. We are unaware of studies other than our current one that document partial and focal BRB compromise specifically in retinal veins. Our results suggest that venous leakage of tracers may become a useful diagnostic test in glaucoma, and a tool in guiding treatment choices. On the other hand, it is possible that different patterns of leakage may occur depending on the magnitude of IOP, type of glaucoma, and/or genetic and environmental background. Although further experiments are needed, it is conceivable that the presence, absence, severity and/or location of vascular leakage may be able to distinguish those whose glaucoma is actively progressing from those in a currently stable state. Another possibility is the use of such tests to differentiate those with ocular hypertension who need treatment because their glaucoma is likely to progress from those with ocular hypertension who are unlikely to progress and do not need treatment. Currently, it is not possible to predictively distinguish these groups. This results in withholding of necessary treatment from some individuals with high IOP but overtreating of others with unnecessary exposure to the potential side effects of these treatments^5,113,114^. An important study recently documented metabolism-related decreases in oxygen consumption and NAD levels in blood cells of glaucoma patients, with lower levels of each associating with a greater rate of glaucoma progression^115^. Along with our current study, this offers promise for valuable, clinically achievable tests to predict glaucoma progression by assessing distinct disease features/mechanisms. We propose that assessing vascular leakage, both with and without assessing NAD and/or metabolism in blood cells, eyes, or other tissues, may provide valuable and rapid tests of treatment efficacy (or of patient compliance). Such test would be important when refining treatment options for individual patients with IOP-lowering or other medications, or even when developing new therapeutic molecules using animal models. BRB leakage could conceivably occur in individuals with normal-tension glaucoma, either for reasons other than IOP, due to a unique sensitivity of their BRB to IOP levels that are not considered elevated, or because higher IOP was missed due to inadequate sampling.

In summary, our study highlights IOP-induced BRB compromise as a previously overlooked yet significant component of glaucoma pathogenesis, paving the way for novel therapeutic interventions and warranting further research.

## Methods

### Mouse strains, breeding, and husbandry

All mice were treated in accordance with the Association for Research in Vision and Ophthalmology’s statement on the use of animals in ophthalmic research. All animal procedures were performed according to the protocols approved by Columbia University’s Institutional Animal Care and Use Committee. DBA/2J mice (strain #000671), DBA/2J-*Gpnmb^+^*/SjJ (strain #007048) and C57BL/6J mice (strain #000664) were purchased from the Jackson Laboratory. Tg(Cdh5-cre/ERT2)1^Rha^ mice^66^ were imported from Dr. Carol Troy’s lab at Columbia University. Ctnnb1^tm1Mmt^ mice^67^ were imported from Dr. Xin Zhang’s lab at Columbia University. Both alleles were backcrossed to DBA/2J for >10 generations to produce congenic mice on the DBA/2J background (all experimental mice were >N10). B6.*Lmx1b^V265D/+^* mice^50^ were from our lab stock. All mice were housed in a 21°C environment with a 14-h light and 10-h dark cycle, fed with a 6% fat diet (PicoLab Rodent Diet 20). Both female and male mice were used for analysis.

### Tamoxifen injection

Tamoxifen (Sigma T5648) solution was prepared at 20 mg/mL in sterile corn oil (Sigma C8267) using a rotator to facilitate dissolution overnight at room temperature, then aliquoted and stored at −80°C. For each mouse, 100 µL of 20 mg/mL tamoxifen in corn oil was administered in 4 intraperitoneal injections spaced 2 days apart.

### Assessment of retinal vascular permeability with tracer

Mice were restrained and intravenously injected with 200 μL 0.02% Hoechst 33342 (Invitrogen, H1399) in 0.9% saline. In some cases, mice were intravenously injected with 100 μg Isolectin GS-IB4 From *Griffonia simplicifolia*, Alexa Fluor 647 conjugate (Thermo Fisher Scientific I32450) in 100 μL 0.9% Saline one-hour prior Hoechst injection to label the retinal vasculature. Fifteen minutes after Hoechst injection, mice were euthanized by cervical dislocation, eyes enucleated, and fixed in 4% paraformaldehyde in 0.1M Phosphate Buffer pH 7.4 for 1 h at room temperature. Retinas were subsequently dissected, flat-mounted onto slides, and imaged with a Leica SP8 laser scanning confocal microscope.

### Immunohistochemistry

Eyes were enucleated and fixed in 4% paraformaldehyde in 0.1M Phosphate Buffer pH 7.4 at room temperature for 1 h. Retinas were dissected and blocked for 4 h at room temperature in blocking solution (Phosphate-buffered saline, 0.5% Triton-X100, 5% donkey serum), and incubated overnight at 4°C with primary antibody in blocking solution. After washing 5 times for 40 min at room temperature with washing solution (Phosphate-buffered saline, 0.5% Triton-X100), retinas were incubated with secondary antibody in blocking solution overnight at 4°C. Retinas were washed for another 5 times for 40 min at room temperature with washing solution, and flat-mounted. The following primary antibodies and lectins were used for this study: Mouse anti-claudin-5, Alexa Fluor 488 conjugate (Thermo Fisher Scientific 352588, 1:500); Rabbit anti-ZO1 (Thermo Fisher Scientific 40-2200, 1:200); Rabbit anti-desmin (Cell Signaling 5332, 1:200); Rabbit anti-NG2 (Sigma AB5320, 1:200); Goat anti-Albumin, FITC conjugate (Thermo Fisher Scientific A90-234F, 1:200); Rabbit anti-Mfsd2a (Cell Signaling 80302, 1:200); Rat anti-CD31 (BD Pharmingen 550274, 1:50); Rabbit anti-RBPMS (Novus NBP2-20112, 1:200) and Isolectin GS-IB_4_ From *Griffonia simplicifolia*, Alexa Fluor 647 conjugate (Thermo Fisher Scientific I32450, 1:50). For each genotype and each antibody, at least 16 eyes were stained for DBA/2J and controls, 8 eyes for *Lmx1b^V265D/+^*, 8 for HAMA and their controls. After staining, retinas were flat-mounted and examined by a Keyence BZ-X810 microscope. All stained eyes were visually analyzed while a subset of them (subset numbers indicated in figure legends) were imaged by a Leica SP8 laser scanning confocal microscope with 63X objective lens for further analysis. Findings agreed between the Keyence and Confocal analyses. Images were stitched with Imaris stitcher 10.0.

### Nerve staining and evaluation of damage

The intracranial segments of the optic nerves were fixed in 0.8% paraformaldehyde; 1.22% glutaraldehyde; and 0.08 M Phosphate Buffer pH 7.4 at 4°C for 12 h, followed by processing and embedding in plastic. The retro-orbital ends of the nerves were cut into 1 μm thick sections and stained with paraphenylenediamine (PPD). PPD stains the myelin sheath of all axons, while specifically darkly staining the axoplasm of damaged axons^116^. At least 3 sections were examined to determine the level of damage for each nerve. The optic nerves were determined to have 3 levels of damage as previously reported and validated against axon counting^12,14^: 1. No or early damage (NOE) - less than 5% of axons were damaged, and there was no gliosis. These nerves do not have glaucoma (no damage by conventional criteria) but are called no or early as some of them have early molecular changes that precede neurodegeneration. 2. Moderate damage (MOD) - had an average of 30% axonal loss and early gliosis. 3. Severe damage (SEV) - had more than 50% axonal loss and damage with prominent gliosis. All nerves were evaluated by at least two masked investigators. In cases where the two investigators did not agree on the damage level a third investigator (also masked) analyzed the nerve and the most assigned damage level was used^117^.

### Image analysis and quantification

Vascular leakage: Hoechst tracer-labeled retinal images were processed with Imaris 10.0 software to detect and quantify vascular leakage. Smoothed surfaces were created based on Hoechst signal (blue wavelength, pseudo colored to red for enhanced visualization in figure) with background subtraction and splitting of touching objects. All created surfaces were then filtered in Imaris to separate endothelial cells from labeled neurons using its inbuilt AI machine-learning tools. In the first round of AI training, several examples of each cell type category were manually assigned. After the next round of AI prediction, several incorrect results were manually corrected and used for the following training round. After 5 rounds of training, the trained model was saved and applied to other images for automated cell-type discrimination. For each new image, additional 1-2 rounds of training were applied. The amount of leakage was quantified by counting Hoechst-positive neuronal nuclei in 0.81mm^2^ regions of the compressed z-stack that were centered over the Y branches of both peripheral veins for each flat-mounted retina. Maximum intensity projections were used to obtain the following stacks: 14 μm z-stacks for GCL, starting from the peripheral veins toward the IPL; 21 μm z-stacks for INL/IPL, starting from the IPL vessels toward the GCL layer; and 21 μm z-stacks for OPL/ONL, starting from the OPL vessels toward the ONL.

RGC quantification: RBPMS-stained RGCs in high resolution images of whole retinal flat mount images were counted by Imaris 10.0 software Spots function. Each individual RGC was marked by a spot according to RBPMS signal and total number of spots were counted.

Tight-junction and mural cell protein: fluorescence signals were quantified in high resolution retinal flat mounts. The “freeform” selection tool in Image J was used to obtain the mean fluorescence value exclusively for the major veins in each retina.

### IOP assessment and induction of high IOP

IOP was assessed using the microneedle method as previously outlined in detail ^118,119^. This prevents overestimation of IOP by noninvasive IOP-measuring instruments due to corneal property changes that can occur in the models used. In brief, mice were anesthetized with an intraperitoneal injection of ketamine (99 mg/kg; Ketlar, Parke-Davis, Paramus, NJ, USA) and xylazine (9 mg/kg; Rompun, Phoenix Pharmaceutical, St Joseph, MO, USA) immediately before the IOP measurement. All IOP measurements were taken during the same time-period each day. We also assessed anterior chamber (AC) depth in all studied eyes as an indicator of exposure to high IOP. All studied ocular hypertensive animals had an obviously enlarged AC, while the space in the AC is tiny in normotensive mice (for representative examples see Fig S7) This ensured that all eyes with genotypes or experimental treatment that increase IOP had experienced high IOP, as increased AC depth is a reliable measure of even modest exposure to high IOP in mice^42,50,57^. When IOP becomes elevated, the measured IOP values of a mouse population spread in each direction as homeostatic regulation and diurnal cycles are perturbed ^120^. This is well established in various mouse models, with some mice having higher IOP during the light-period of the day and others during the dark cycle^119,121,122^. Thus, among eyes with abnormal IOP exposure, high IOP is not detected in every eye at every measurement time and AC depth serves as a reliable surrogate ^42,50,57^. Increased AC depth was never observed in control normotensive mice. For the experimentally induced model, IOP was elevated by polymerizing a hydrogel resistor in the path of aqueous humor drainage as published^51^ with some modifications in C57BL/6J mice at 3 months of age. Instead of penetrating the cornea with a 32 G needle, a small incision was made with a very sharp sapphire knife to prevent corneal damage from the surgery. One μL of HAMA was delivered into the angle using a 100-micron OD glass microneedle. Using this method, IOP becomes promptly elevated, and the higher values are sustained for about 4 weeks.

### Western blots

Retinas were dissected in ice cold phosphate-buffered saline, homogenized in lysis buffer (150 mmol/L NaCl, 50 mmol/L Tris HCl pH 8.0, 1 mmol/L CaCl_2_, 1% NP40, 0.1% N-dodecyl-β-D-maltoside, 3 mmol/L MgCl_2_, and DNAseI 80 units/mL) with protease inhibitor (Roche, cOmplete 4693116001) and incubated on ice for 30min. Retinal lysates were then spun down at 14000 rpm for 5 min. 160 μL supernatants were mixed with 40 μL 1M DTT and 200 μL Laemmli sample buffer (Bio-Rad, 1610737), incubated at 40°C for 10 min, run on 4%-15% polyacrylamide gel (Bio-Rad, 4561084), and transferred to PVDF membranes. The following primary antibodies were used for this study: Rabbit anti-claudin-5 (Cell Signaling 49564S, 1:1000); Rabbit anti-ZO1 (Thermo Fisher Scientific 40-2200, 1:4000); Rabbit anti-occludin (Thermo Fisher Scientific 40-6100, 1:5000); Mouse anti-β-actin, HRP conjugate (Thermo Fisher Scientific MA5-15739-HRP, 1:20000). Chemical signals were imaged by iBright system (Invitrogen). Signals were quantified with ImageJ gel plugin and normalized by β-actin.

### Statistical analysis

Fluorescence intensity for immunofluorescence and Western blot signals (**Figure 2**) were compared using Welch’s t-test for each marker. In the β-catenin stabilization experiments (**Figure 5**), numbers of leakage labeled RGCs (Hoechst-positive neurons per mm^2^) and total RGC counts (RPBMS) were compared using Welch’s t-test. Fisher’s Exact test was used to compare the degree of nerve damage between control and stabilized β-catenin-expressing groups. Control and stabilized β-catenin-expressing IOPs were compared using Welch’s t-test.

## Acknowledgements

The authors would like to thank Dr. Chenying Guo and Dr. Ganesh Prasanna for their help when establishing the HAMA injection method, Pete Finger for optic nerve sectioning and PPD staining, Dr. Matthew Gastinger for image analysis guidance with Imaris, Dr. Carol Troy for sharing the Tg(Cdh5-cre/ERT2)1^Rha^ mice, Dr. Xin Zhang for sharing the Ctnnb1^tm1Mmt^ mice, Dr. David Silver and Dr. Chenghua Gu for sharing their MFSD2A antibodies at the early stage of this project before it was commercially available, Devanshi Ragha for her assistance, and Institute of Comparative Medicine staff at Columbia University for mouse husbandry and assistance. The authors also thank Dr. Richard Libby, Dr. Krishnakumar Kizhatil, Dr. Michael Elliott, and members of the John lab for helpful comments, advice and assistance at various stages of the project.

---

**Supplementary Figure 1.**
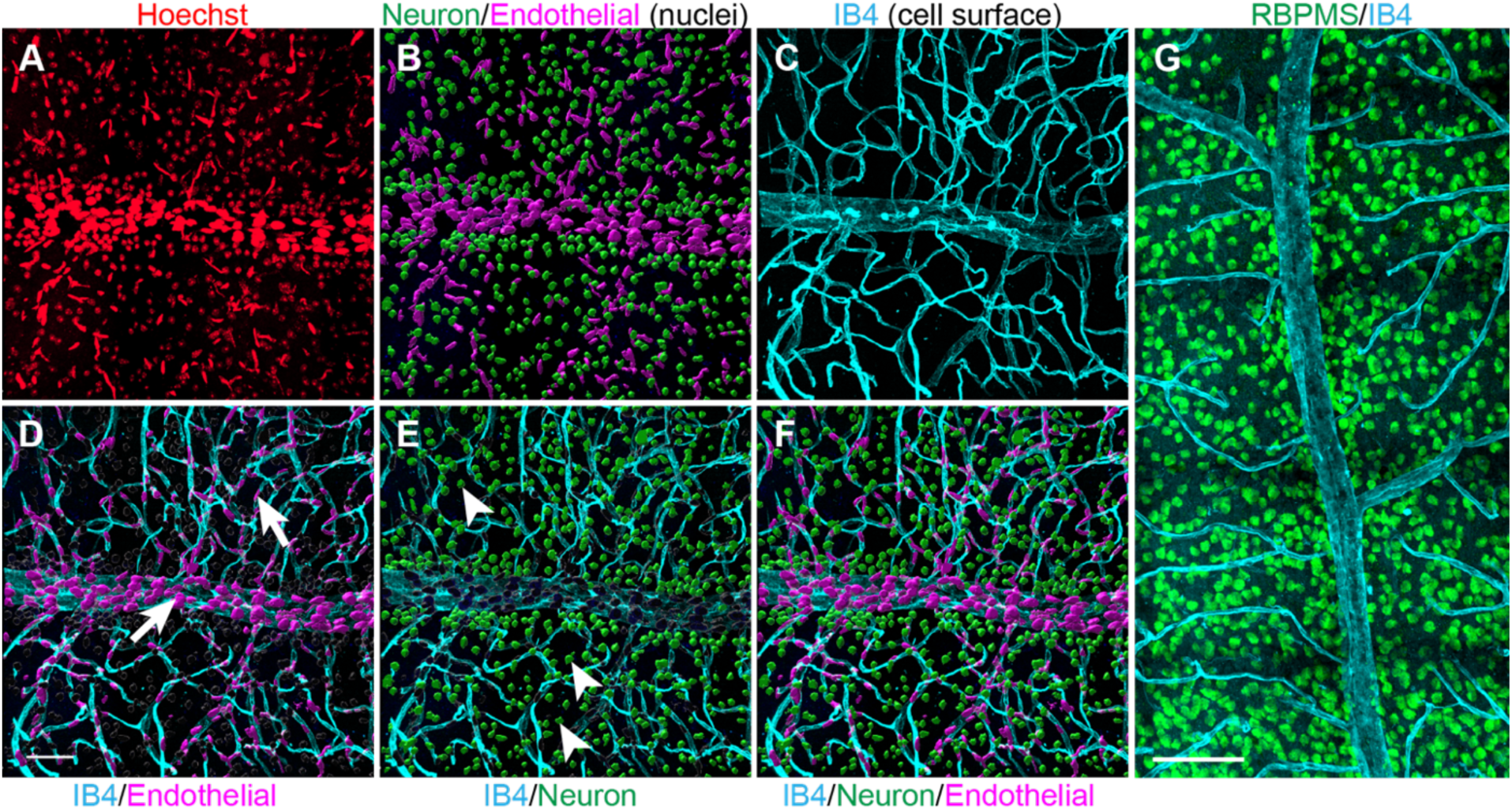
Differentiating neural vs. endothelial cell nuclei for quantification. Representative images from retinal flat mounts. **A.** Whole flat mounts from 4 Hoechst-administered mice were used to train a machine-learning model in Imaris to identify and differentiate between neural and endothelial cell nuclei. **B.** AI-filtered images showing neural (green), and endothelial (magenta) cell nuclei. **C.** IB4 (cyan), a marker for endothelial cells, was used for comparison/confirmation. **D-F.** The nuclei that the AI identified as endothelial (arrows, D) co-localized with IB4 staining, whereas the nuclei identified as neural (arrowheads, E) did not. The distribution, of neuronal nuclei matches the distribution of neurons (see G) demonstrating the accuracy of our model. F is a merge of the same retina shown in A-E. **G.** RBPMS-stained retinal ganglion cells (neurons). Scale bar: 50 μm in A-F, 100 μm in G.

**Supplementary Figure 2.**
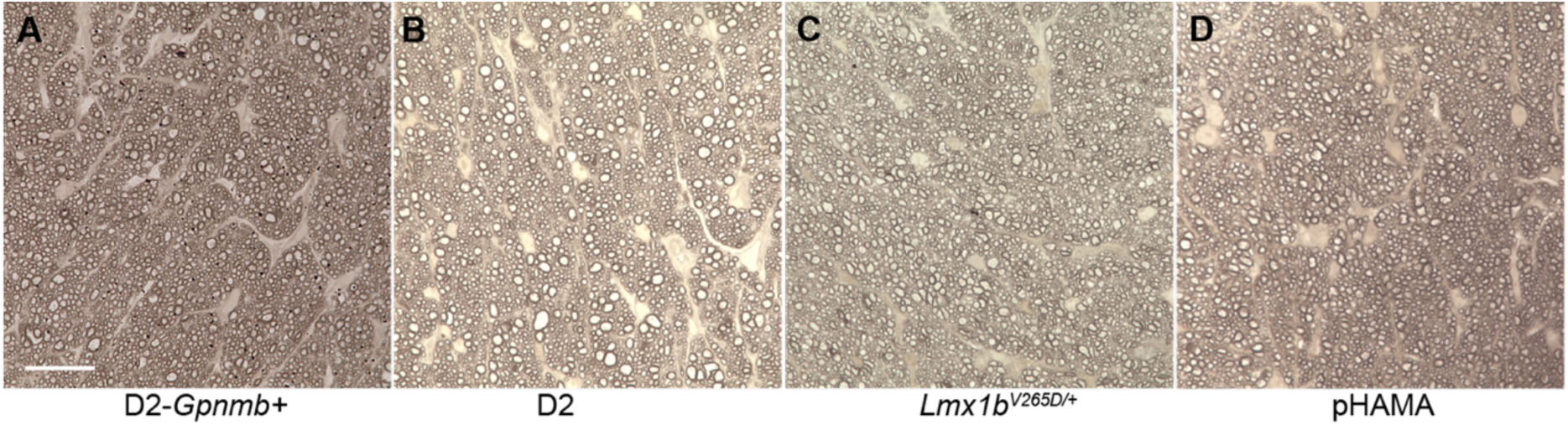
Optic nerve integrity in the presence of BRB leakage. **A-D**) Representative images from PPD-stained optic nerve cross sections. PPD differentially stains damaged axons, allowing a very sensitive detection of optic nerve damage (Methods). The optic nerve images in B-D are from the same mice as in Figures 1A (B), S4A (C), and S4C (D), where leakage was demonstrated. At these ages, they all have healthy optic nerves that are indistinguishable from controls (A). Together with the other figures, this demonstrates that BRB compromise and vascular leakage precede neurodegeneration in all three glaucoma models. pHAMA= polymerized HAMA. Scale bar: 20 μm.

**Supplementary Figure 3.**
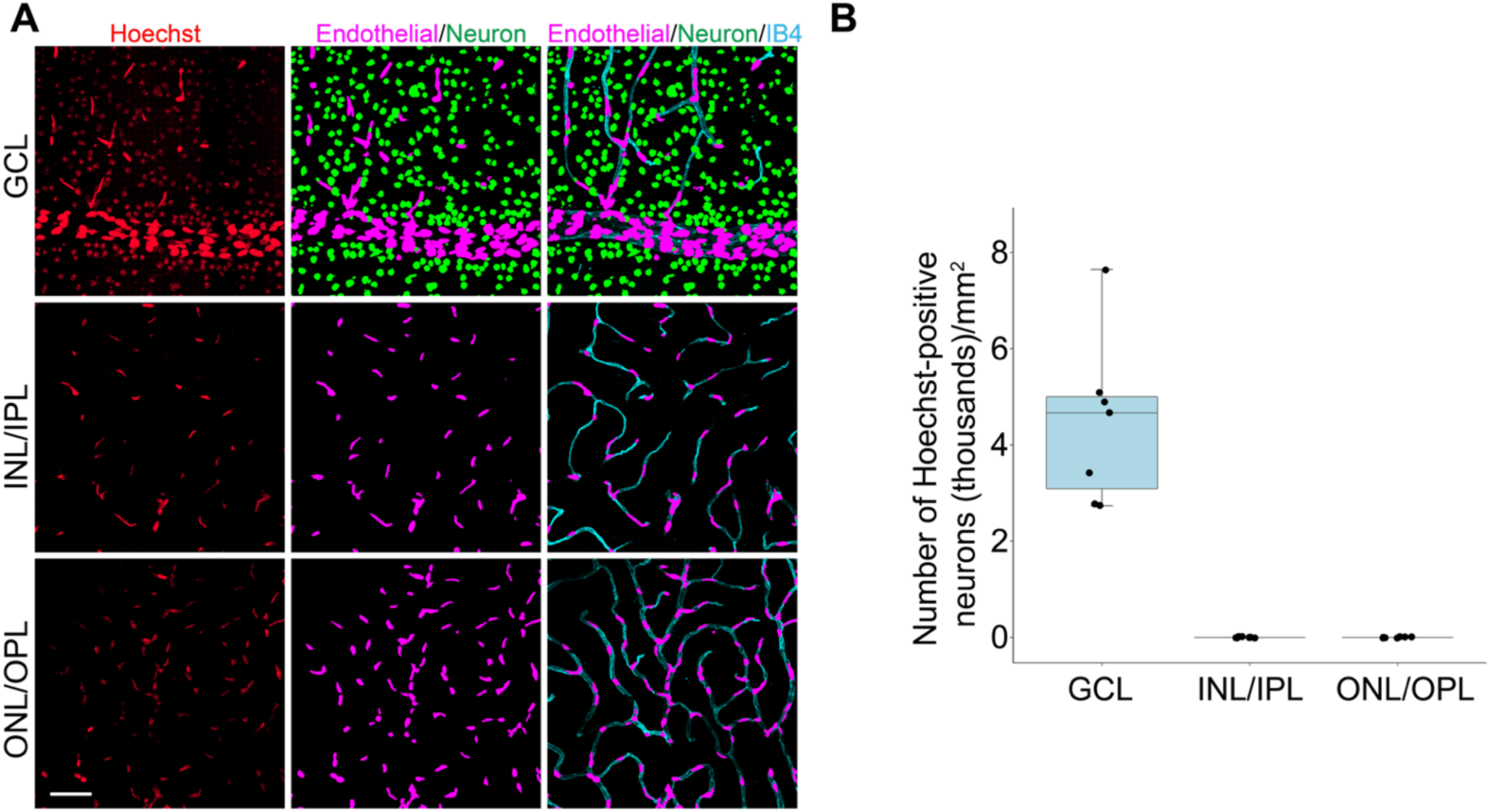
BRB leakage is restricted to the ganglion cell layer of DBA/2J mice at late disease stages. **A.** Optical sections of retinal flat mounts at different tissue depths. IB4 staining (cyan) is a marker of endothelial cells and co-localizes with Hoechst-stained endothelial cell nuclei identified by machine learning (in magenta). Even at this advanced age (14 months), with longer pressure/disease exposure, leakage only labels neurons (green) in the ganglion cell layer (GCL). Inner and outer nuclear layer (INL/ONL) neurons are not labeled indicating that leakage does not occur from vessels of the inner or outer plexiform layers (IPL/OPL). **B.** Quantification of leakage in different retinal layers. Scale bar: 50 μm. n = 20 visually analyzed, n = 7 quantified.

**Supplementary Figure 4.**
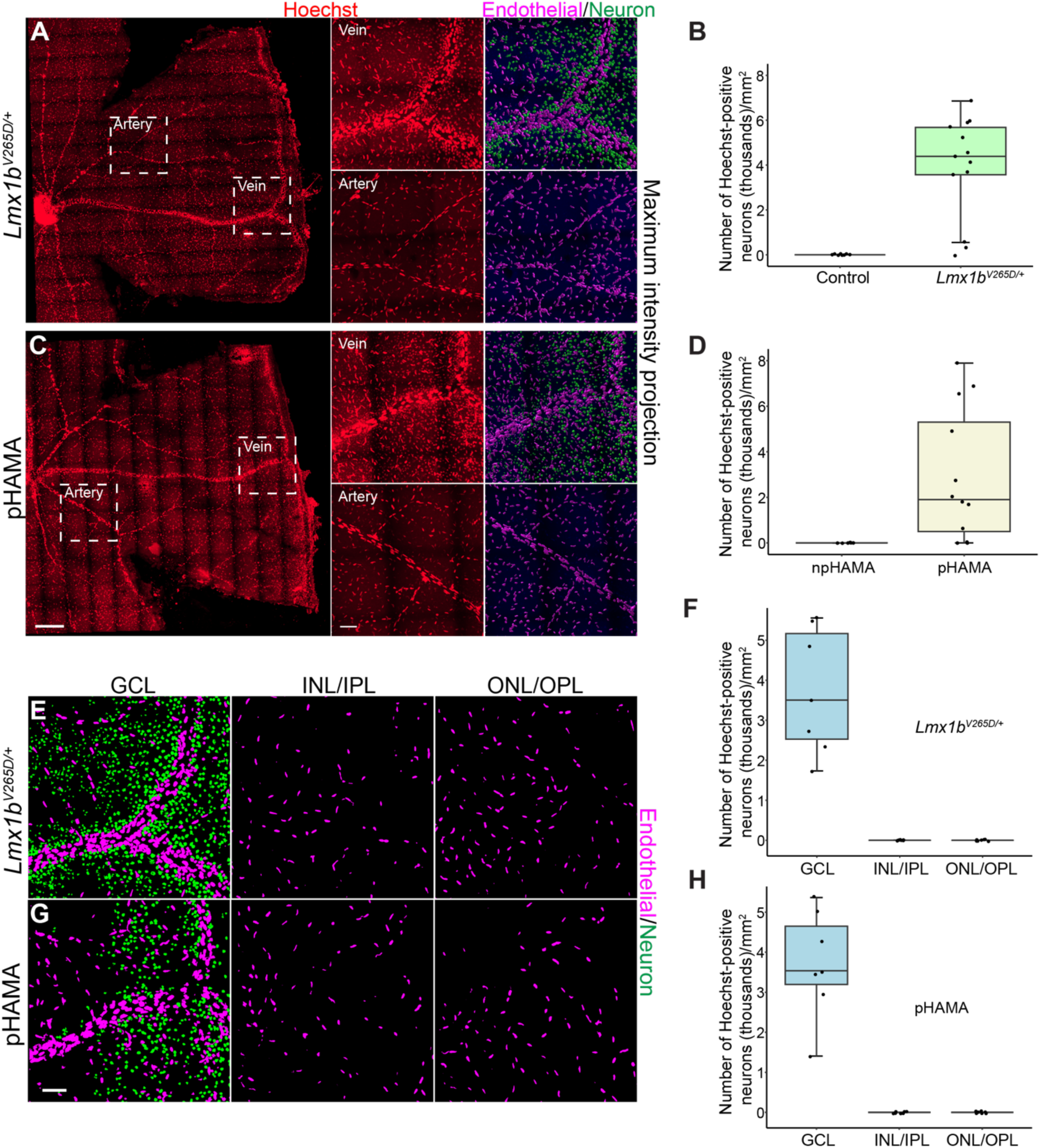
IOP-induced BRB breakdown in additional mouse models. A-B, Detection of extravascular Hoechst (red) indicates BRB compromise in another genetic model with increased IOP prior to glaucoma. Leakage was only detected from peripheral veins (not arteries or capillaries). B6.*Lmx1b^V265D/+^*: 5.5-month-old eyes with confirmed exposure to high IOP by deepened anterior chambers (ACs) but no glaucomatous damage (see Figs: S5 optic nerve and S6 IOP distribution this model). *Lmx1b^+/+^*, n = 8; *Lmx1b^V265D/+^*, n = 13. **C-D.** We observed the same pattern of vascular leakage following experimental elevation of IOP by polymerization of HAMA in the drainage angle (pHAMA). Four-month-old C57BL/6J eyes one month after polymerization of HAMA. Exposure to high IOP was confirmed by deep ACs (see Fig. S7) but there was no glaucomatous damage (see Fig. S2). HAMA rapidly raises IOP (see Fig. S7). npHAMA, n = 7; pHAMA, n = 12. Above are representative images of retinal flat mounts including arteries and veins (scale bar: 300 μm) with regions in white boxes magnified to the right (scale bar: 50 μm). Endothelial (magenta) and neuronal (green) nuclei were color-coded in Imaris as described in Figure S1. **E-H.** Leakage was restricted to the veins abutting the ganglion cell layer (GCL) with leaked tracer labeling neurons in the ganglion cell layer but not in any retinal layer from the INL to ONL. *Lmx1b^V265D/+^*, n = 7; pHAMA, n = 7.

**Supplementary Figure 5.**
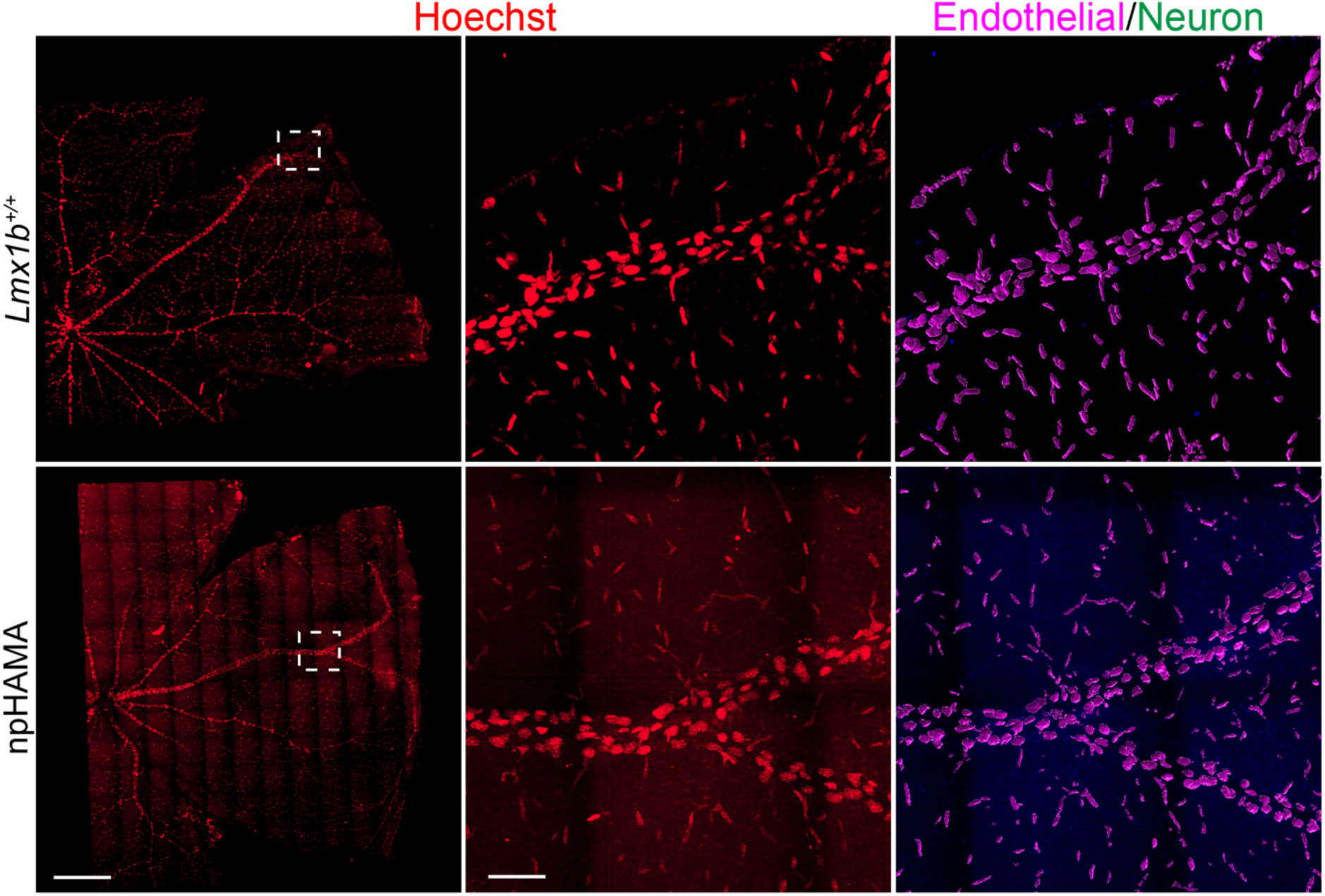
No BRB leakage in age-matched normotensive control mice. Retinal flat mounts from the ocular normotensive control mice (*Lmx1b*^+/+^ or npHAMA) that were administered Hoechst. White-boxed regions are magnified to the right. No neurons were stained by Hoechst as the BRB was intact. Scale bar left panels: 300 μm; mid and right panels: 50 μm.

**Supplementary Figure 6.**
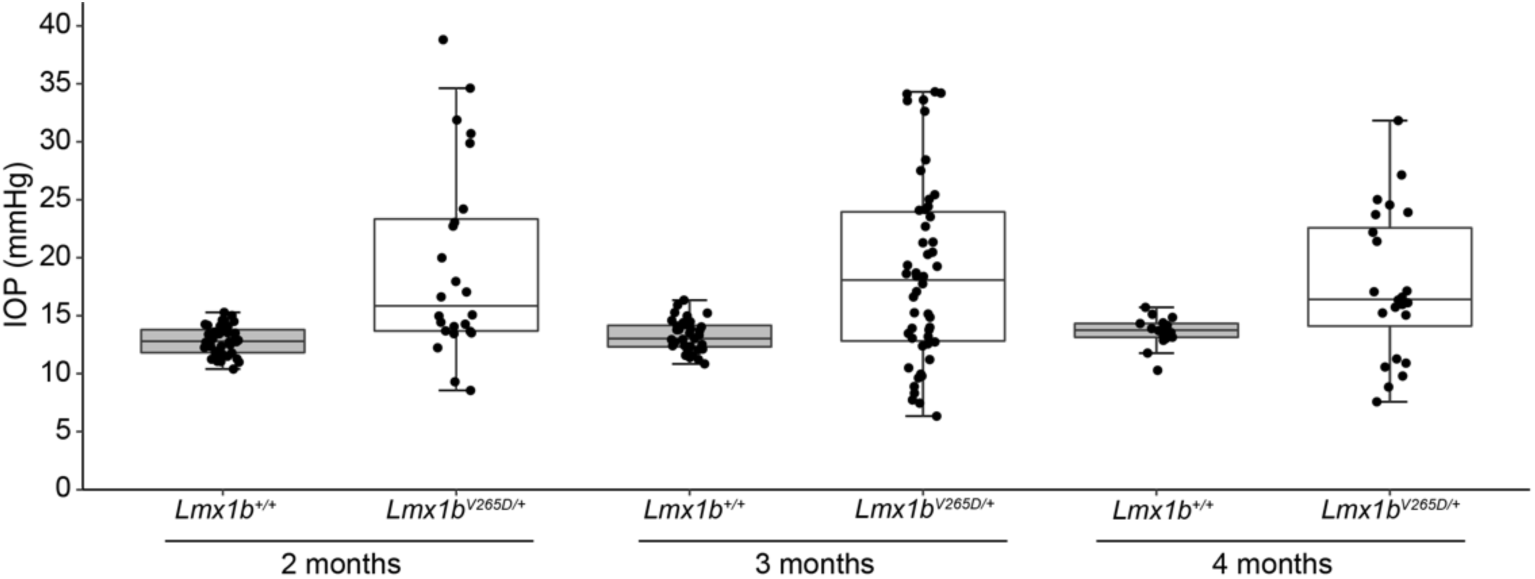
IOP distribution in B6.*Lmx1b^V265D^* mice. Boxplots of daylight time IOP in B6.*Lmx1b^V265D/+^* mice and control mice with age. IOP elevation is first seen at 2 months of age in B6.*Lmx1b^V265D/+^* mice but not in control mice. As with other mouse models of glaucoma, high IOP/IOP elevating pathology disturbs ocular homeostasis. At ages when IOP elevation is detected in these models, the IOP spreads in both directions in a population of mice (see ref 50). IOP is typically higher in more mice during the dark period of the day and the IOP of individual eyes varies with time of measurement (see refs. 119, 121, 122). As high IOP causes corneal stretching and anterior chamber deepening, a deep anterior chamber is a reliable indicator that any eye has experienced high IOP. 2 months: *Lmx1b^+/+^*, n=44; *Lmx1b^V265D/+^*, n = 24; 3 months: *Lmx1b^+/+^*, n = 35; *Lmx1b^V265D/+^*, n = 50; 4 months: *Lmx1b^+/+^*, n=17; *Lmx1b^V265D/+^*, n = 24.

**Supplementary Figure 7.**
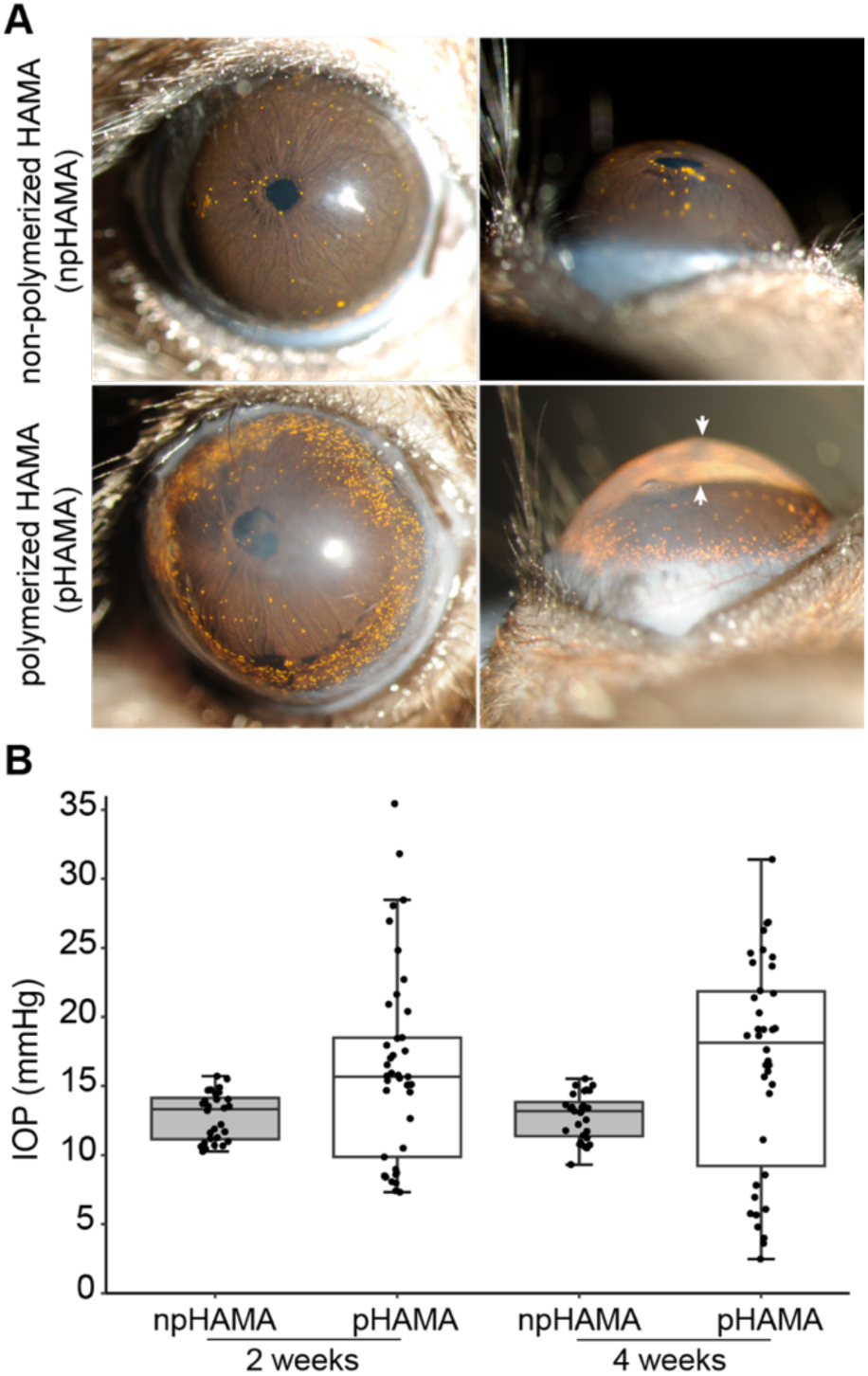
HAMA model of experimentally induced IOP. **A.** Slit-lamp images show anterior chamber deepening in eyes with high IOP induced by polymerization of HAMA (pHAMA) but not in normotensive control eyes with non-polymerized HAMA (npHAMA). All images 2 weeks after injection of HAMA. npHAMA eyes underwent the same procedure as HAMA eyes except that HAMA was not polymerized. As HAMA is transparent, it was mixed with microbeads to enable visualization of the polymerized hydrogel ring around the ocular periphery and angle. The pHAMA eye has an obviously deepened anterior chamber (arrows) to a similar extent as used for confirmation of exposure to high IOP throughout our experiments. **B.** IOP distribution demonstrating IOP elevation in pHAMA but not control eyes. 2 weeks: npHAMA, n=28; pHAMA, n=41; 4 weeks: npHAMA, n=28; pHAMA, n=38.

**Supplementary Figure 8.**
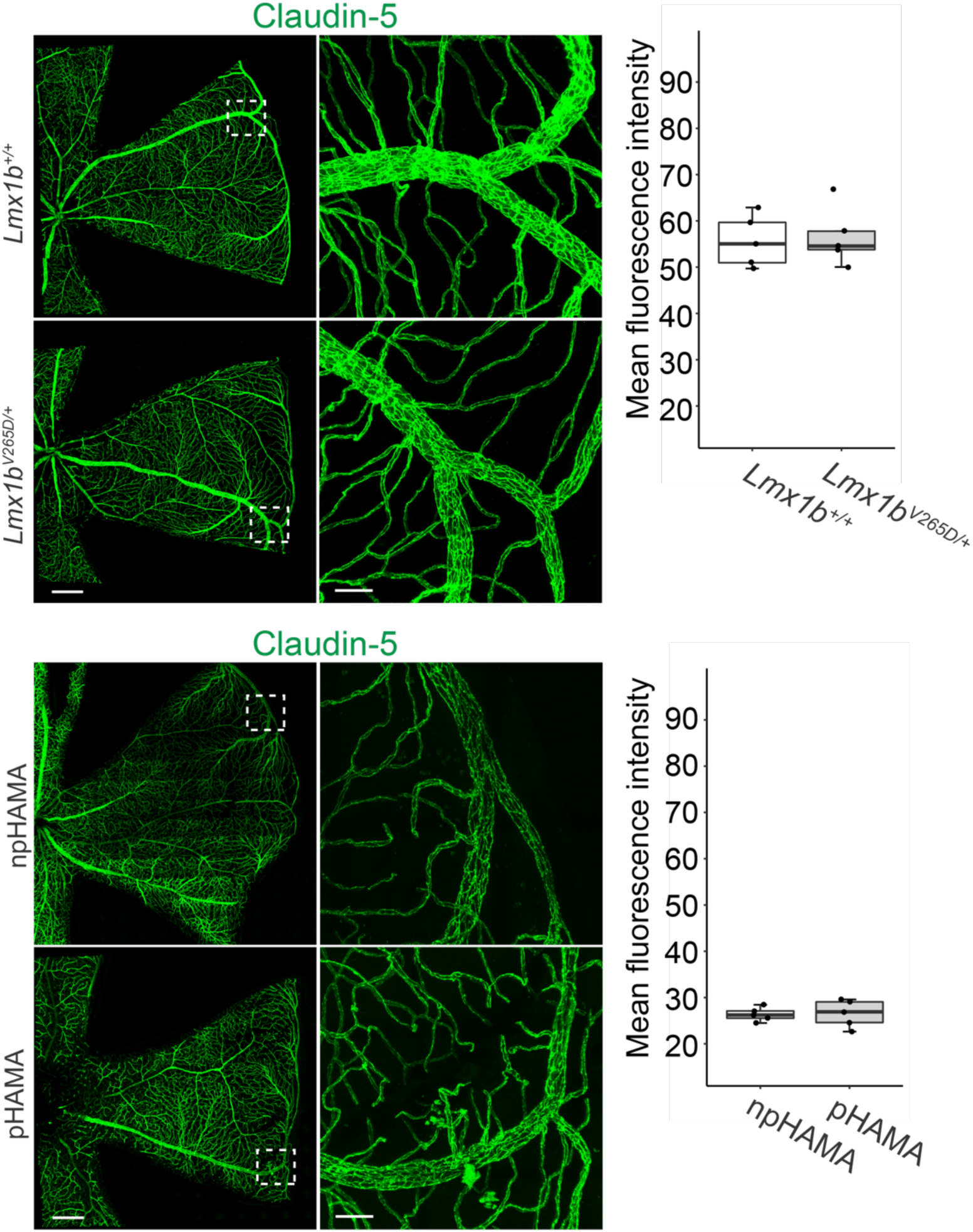
Normal tight junction marker in hypertensive B6.*Lmx1b^V265D/+^* and HAMA eyes. Representative images of staining with claudin-5 in retinal flat mounts of B6.*Lmx1b^V265D/+^*mice (6.5-month-old) and ocular hypertensive pHAMA mice (4-month-old, 1 month after injection) along with their normotensive controls. No difference in this tight junction marker was detected (n=5 veins, from 5 eyes each group, *P* = 0.8087 (top), *P* = 0.8982 (bottom)). Paired images with boxed regions at higher magnification in the adjacent images to the right. Scale bars: left images, 300 μm; right images, 50 μm.

**Supplementary Figure 9:**
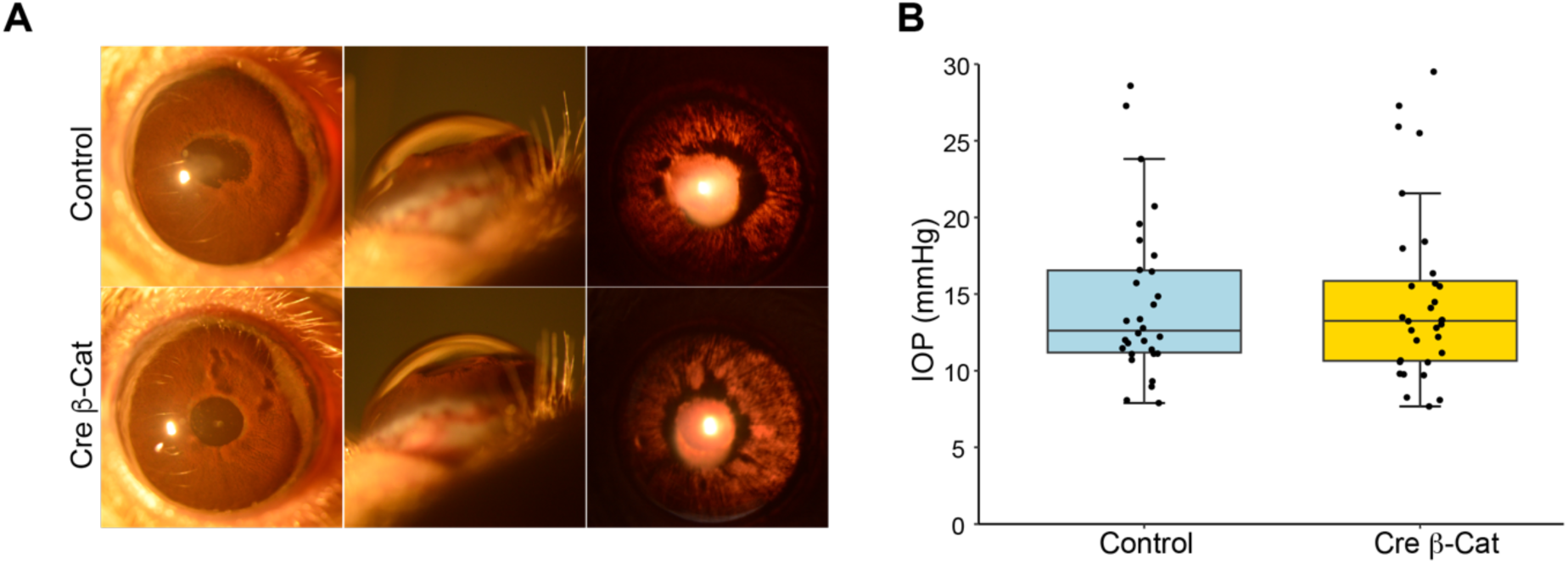
Stabilization of β-catenin does not affect IOP elevating iris disease or IOP itself. A. Representative slit-lamp images of D2. *Ctnnb1^flex3^*^/+^ (Control) and D2.*Ctnnb1^flex3^*^/+^; Cdh5-CreERT2 (Cre β-Cat) mice eyes (8-month-old). A pigment-dispersing iris disease elevates IOP in D2 mice. The anterior segment phenotypes did not differ in timing or severity between genotypes. Left panels: obvious iris atrophy and pigment dispersion. Middle panels: similarly deepened anterior chambers (a consequence of IOP elevation) A space between iris and cornea is hardly detectable in ocular normotensive mice (see Fig. S7). Right panels: similar transillumination defects where light passes through the depigmented iris. See Fig. S7 for an unaffected eye. **B.** No difference in IOP was observed between Control and Cre β-Cat mice (Welch’s t-test, *P* = 0.89) at 8-month-old. Control; n = 30, Cre β-Cat; n = 28.

**Supplementary Figure 10.**
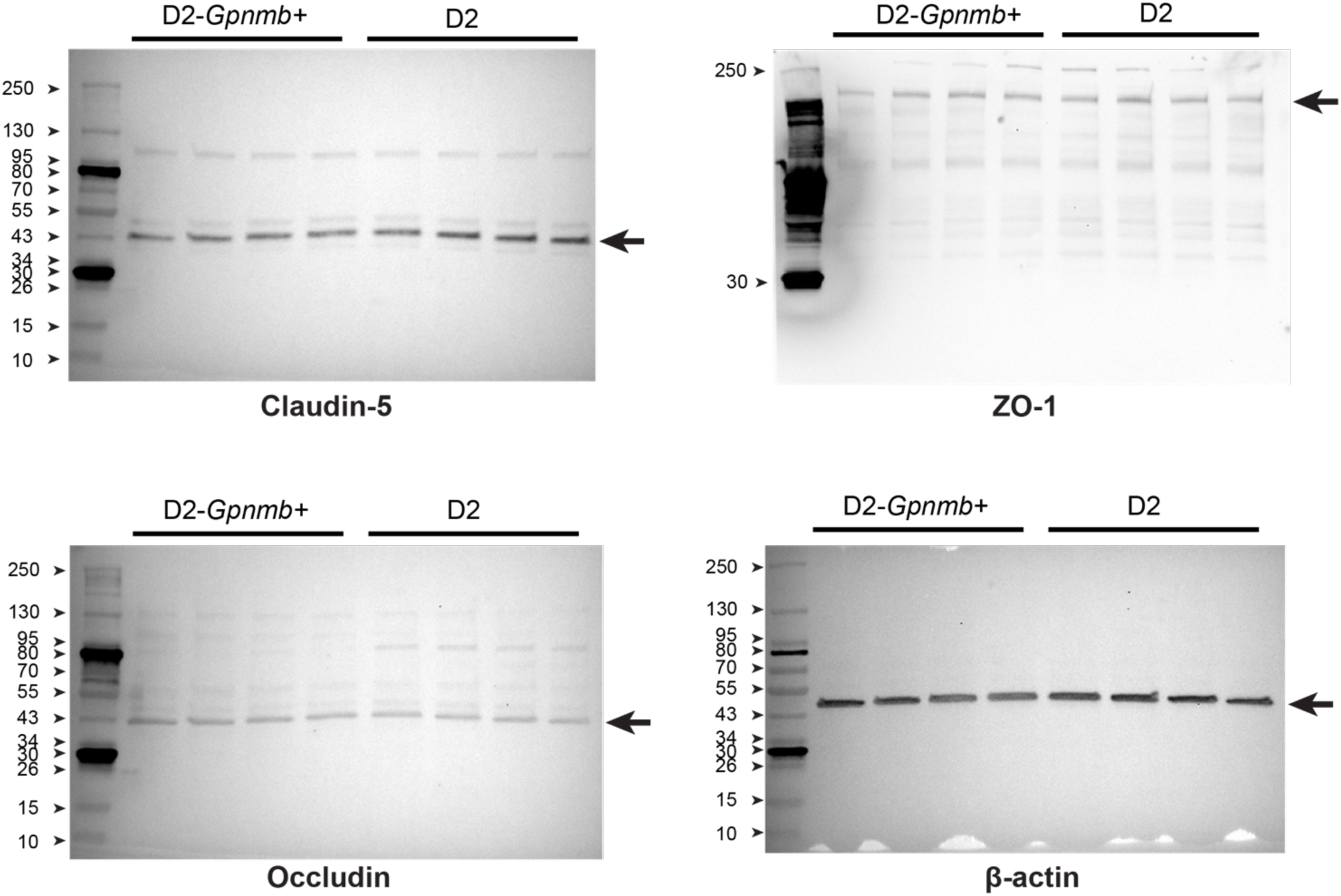
Images of original Western blots from Figure 2. Position of target protein indicated by arrow.

